# ARNy Plotter: A Comprehensive Web Server for RNA Ensemble Structural Analysis and Visualization

**DOI:** 10.64898/2025.12.24.696344

**Authors:** Louis Meuret, Julien Rey, Samuela Pasquali

## Abstract

Analyzing RNA structural ensembles, whether derived from molecular dynamics, enhanced sampling techniques, or experimental data, poses significant challenges due to the intrinsic flexibility and diverse conformational landscapes of RNA. We present ARNy Plotter, a web-based platform designed for comprehensive analysis and visualization of RNA structural ensembles. ARNy Plotter integrates multiple state-of-the-art methods into a unified, user-friendly environment, enabling intuitive examination of RMSD and eRMSD distributions, torsion-angle variability, dynamic secondary-structure patterns, contact-frequency networks, and multidimensional conformational landscapes. By leveraging MDAnalysis, Barnaba, and foRNA, the platform supports all major trajectory and ensemble formats while offering real-time 3D visualization via MolStar. Unlike existing tools that focus primarily on static structures or provide limited dynamic analysis, ARNy Plotter allows in-depth exploration of conformational transitions and population distributions within RNA ensembles, and facilitates result sharing without distributing full datasets. This platform addresses the growing need for accessible, comprehensive, and interactive tools for RNA structural biology. ARNy Plotter is freely available at https://arny-plotter.rpbs.univ-paris-diderot.fr/, requires no installation or registration, and can also be run offline with a local webserver, including an API for automated figure generation.

## 1 Introduction

RNA molecules are now recognized as central regulators of cellular processes, extending far beyond their traditional role as intermediates in protein synthesis. They function as regulatory elements, catalytic enzymes (ribozymes), and structural scaffolds, with over 60% of the human genome transcribed into non-coding RNAs [1]. This functional variability arises from RNA’s ability to adopt complex three-dimensional architectures through networks of base pairing, stacking interactions, and tertiary contacts. Understanding RNA function therefore requires characterization of both stable structures and time-dependent conformational changes. Many RNA-mediated mechanisms rely on large-scale structural rearrangements occurring on microsecond-to-millisecond timescales [2, 3], particularly in regulatory RNAs where conformational switching governs activity. For example, riboswitches modulate gene expression through ligand-induced structural transitions, while ribozymes depend on specific rearrangements to achieve catalytic activity [4, 5].

Computational approaches such as Molecular Dynamics (MD) and Path Sampling (PS) have become indispensable for investigating RNA conformational dynamics at atomic resolution [6, 7]. These methods generate detailed trajectories that capture the temporal evolution of atomic coordinates, offering critical insights into RNA folding, unfolding, and functionally relevant conformational transitions. Contemporary MD simulations routinely access nanosecond- to microsecond-timescale dynamics, encompassing many biologically relevant processes, and recent advances have enabled millisecond-scale simulations of large RNA systems [8]. PS methods enable systematic exploration of the conformational energy landscape, revealing alternative metastable states and identifying the lowest-energy transition pathways between them [9]. Such simulations can produce tens of thousands of structures corresponding to local minima and transition states. Because the wealth of information generated by MD trajectories or PS energy landscapes, specialized analytical approaches are needed for a meaningful interpretation. Moreover, the unique structural features of RNA, including diverse base pairing patterns, flexible backbone conformations, and complex tertiary interactions, demand specific analysis methods that differ substantially from those developed for protein systems [10].

The existing ecosystem of RNA structural analysis tools exhibits significant fragmentation, with most solutions addressing specific aspects of RNA analysis rather than providing comprehensive frameworks.

### Static Structure Analysis Tools

Several established software focus primarily on static structure visualization and analysis. VARNA (Visualization Applet for RNA) provides interactive 2D secondary structure diagrams [11], while RNAView generates automated secondary structure annotations from 3D coordinates [12]. MC-Annotate offers detailed base pair classification and geometric analysis [13], and RNApdbee provides annotations from 3D structures[14]. However, these tools are limited to single-structure analysis and cannot handle trajectory data.

### General-Purpose Molecular Viewers

For dynamic analysis, researchers usually rely on general purpose molecular visualization software such as VMD (Visual Molecular Dynamics) [15] or PyMOL [16]. Although these tools provide powerful visualization capabilities, they require substantial expertise for RNA-specific analyses and lack integrated analytical workflows tailored to RNA structural features.

### Specialized Analysis Libraries

Specialized trajectory analysis often requires custom scripting using computational libraries such as MDAnalysis [17] or MDTraj [18], which is a significant barrier for researchers without extensive programming backgrounds. These libraries, while powerful, require computational expertise and lack user-friendly interfaces, limiting their adoption in the broader research community.

**Web-Based Solutions** for RNA trajectory analysis remain particularly scarce. SimRNAweb2.0 represents one of the few available platforms, focusing on coarse-grained RNA folding simulations [19]. However, this tool is specifically designed for its own simulation method and does not provide comprehensive analysis capabilities for arbitrary MD trajectories generated using different force fields or simulation packages.

To overcome these limitations and provide the RNA community with a comprehensive and accessible platform for structural ensemble analysis, we developed ARNy Plotter. ARNy Plotter streamlines RNA ensemble analysis by integrating multiple complementary analytical approaches into a unified, web-based interface that requires no software installation or programming expertise. The platform leverages established computational frameworks including MDAnalysis for trajectory processing, Barnaba for RNA-specific structural analysis [20], and foRNA (force-directed RNA) for interactive secondary structure visualization [21]. This integration creates a powerful environment that combines the robustness of established algorithms with the accessibility of modern web technologies. ARNy Plotter supports all major molecular structure and trajectory formats, ensuring compatibility with trajectories generated using diverse simulation packages and methodologies. It can be used to analyze any ensemble of RNA data. This includes also data from experiments such as NMR, where one can easily compare the key features of the multiple structures proposed by the experiment.

ARNy Plotter is born as a web interface, but it is also possible to host the platform locally; the required packages are limited, the technology is straightforward, and API and Python interfaces to the platform are provided for easy integration into existing workflows.

## 2 Design and Implementation

ARNy Plotter provides a carefully designed user interface that guides researchers through the analytical workflow without requiring computational expertise. The interface is organized into four primary panels (Figure 1).

**Figure 1:**
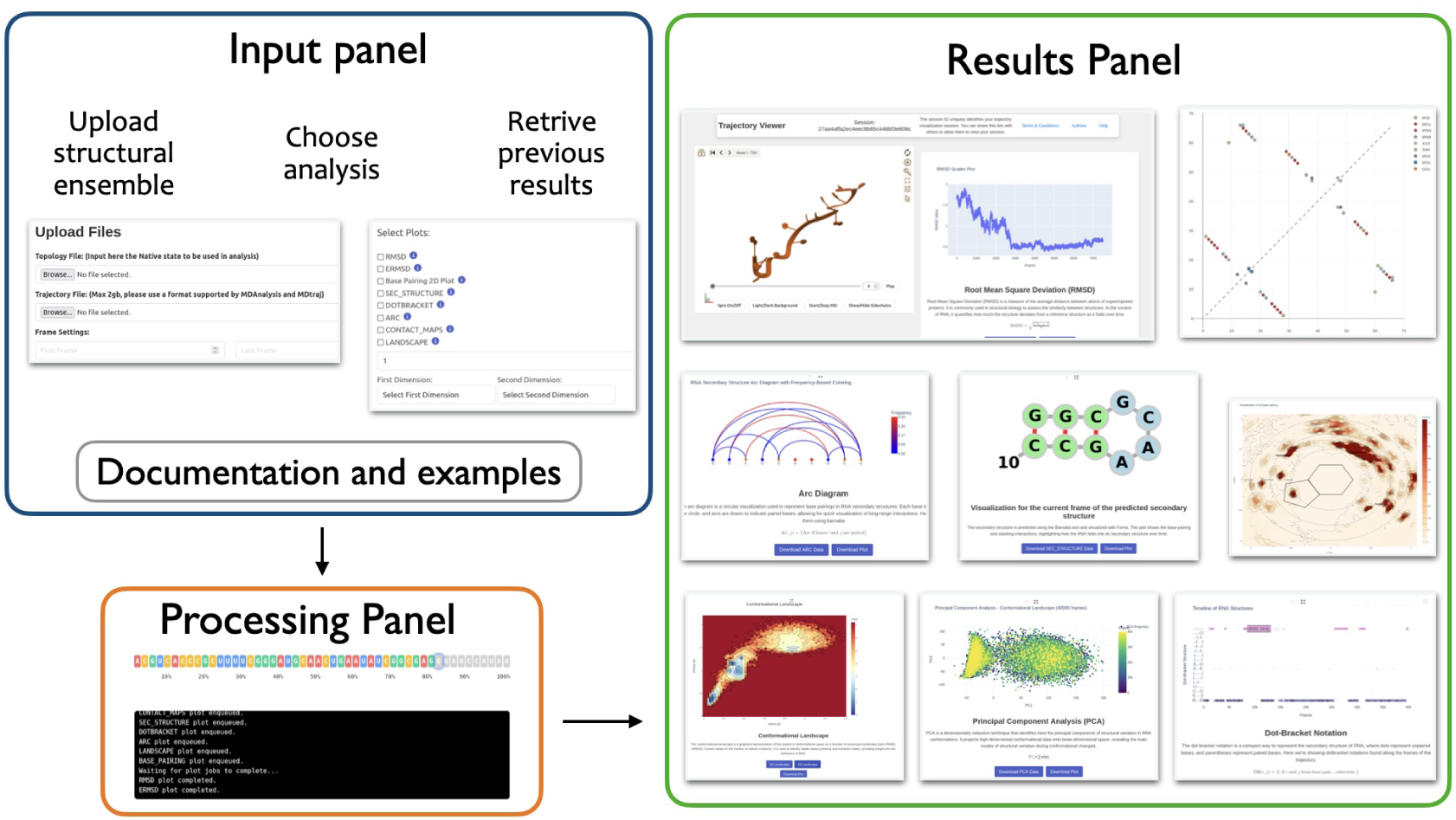
Overview of the ARNy Plotter webserver interface. The input and processing panels handle data upload, analysis selection, and job execution, while the results panel showcases examples of plots generated from structural ensemble analyses.

From the **Input Panel** users can upload the structural ensemble, specify analysis parameters, and select desired visualization outputs. The interface provides clear guidance on file format requirements and parameter selection. It is also possible to input a Session ID, allowing users to retrieve previous calculations from the website or share their session with another user. In the **Processing Panel**, upon job submission, a terminal-style interface displays real-time progress updates for each selected analysis component. Progress bars and status messages keep users informed about computational progress. A button is also available providing a link to the future result page, allowing the user to come back later to visualize the results. The **Results Panel** displays completed analyses in a structured layout that supports easy comparison across multiple visualization types. All results are rendered using responsive design principles. The **Documentation Panel** is accessible from the Input and Results Panels and offers a variety of support resources, including tutorials, parameter explanations, and troubleshooting guides. Each plot in the input panel also includes dedicated buttons that link directly to the relevant sections of the documentation, allowing users to quickly access the information associated with that plot.

### Input Data Processing and Format Support

Input RNA structural ensembles in ARNy Plotter are to be entered as “trajectories”, i.e. successively linked frames of a same molecule. The software provides comprehensive support for molecular structure and trajectory formats through integration with MDAnalysis, enabling processing of trajectories generated using virtually all contemporary formats, including GROMACS (TRR, XTC, GRO formats), AMBER (NetCDF, DCD formats), CHARMM (DCD, PSF formats), and NAMD (DCD, PDB formats). The input interface requires two primary data components: a topology file and a trajectory file. The **Topology File** provides essential molecular structure information, including atomic connectivity, residue definitions, and system topology. Supported formats include PDB, PSF, TOP, GRO, and other MDAnalysis-compatible topology formats. The topology file currently serves as the structural reference for all subsequent analyses, for example, RMSD or eRMSD. The **Trajectory File** contains the ordered series of atomic coordinates. Supported formats include TRR, XTC, DCD, NetCDF, and other standard trajectory formats.

Since for large systems and large conformational ensembles analysis can be computationally expensive, the platform provides the option of trajectory sub-sampling, allowing users to specify the initial frame, final frame, and stride (sampling frequency). This functionality enables focused analysis of specific portions of the trajectory when necessary.

### Ensemble’s analysis and visualization

Once the trajectory is uploaded and processed according to the initial input, the result panel exhibit the analysis results, featuring the trajectory visualization on the left hand side, and plots of the analysis on the right hand side.

#### Interactive 3D Trajectory

The left piece of the results interface features an embedded MolStar molecular viewer that provides 3D visualization of RNA trajectories. MolStar represents the state-of-the-art in web-based molecular visualization, offering performance and functionality close to desktop applications. The trajectory viewer includes playback controls: Frame slider for direct navigation to specific time points; Step-by-step navigation buttons for more detailed frame analysis; Continuous playback functionality; Synchronization with 2D analytical plots for coordinated analysis. This integration enables researchers to correlate visual structural changes with quantitative metrics, facilitating deeper understanding of structure-function relationships.

**Root Mean Square Deviation (RMSD)** calculations provide fundamental metrics for assessing conformational similarity between frames and reference structure. Additionally, ARNy Plotter implements the escore-RMSD (eRMSD) metric, which provides RNA-specific structural similarity assessment focused on functionally relevant geometric features rather than raw atomic coordinates. The eRMSD metric emphasizes base pairing geometries and stacking interactions, providing more biologically meaningful similarity measurements for RNA structures [20].

Interactive scatter plots display structural similarity metrics across the structural ensemble. Hovering interactions reveal precise values and corresponding frame numbers, clicking on one of the point show the corresponding frame in the molstar viewer.

#### Torsion Angles

RNA backbone flexibility is characterized by seven primary torsion angles that define the conformational state of each nucleotide residue. ARNy Plotter provides torsion angle analysis capabilities for a chosen nucleotide, calculating and visualizing the temporal evolution of these structural parameters throughout the ensemble. The analyzed torsion angles are: *α* (O3’*_i−_*_1_ - P*_i_* - O5’*_i_* - C5’*_i_*), *β* (P*_i_* - O5’*_i_* - C5’*_i_* - C4’*_i_*), *γ* (O5’*_i_* - C5’*_i_* - C4’*_i_* - C3’*_i_*), *δ* (C5’*_i_* - C4’*_i_* - C3’*_i_* - O3’*_i_*), *ɛ* (C4’*_i_* - C3’*_i_* - O3’*_i_* - P*_i_*_+1_), *ζ* (C3’*_i_* - O3’*_i_* - P*_i_*_+1_ - O5’*_i_*_+1_), *χ* (O4’*_i_* - C1’*_i_* - N9*_i_*/N1*_i_* - C4*_i_*/C2*_i_*). These calculations enable identification of conformational preferences, backbone flexibility regions, and transitions between distinct torsional states that may correlate with functional conformational changes. Users can visualize torsion values either as a scatter plot or as a histogram.

**Secondary Structures** analysis is a cornerstone of RNA structural characterization. Leveraging Barnaba structural annotations, ARNy Plotter provides three complementary visualization modalities for comprehensive secondary-structure analysis: (i) interactive two-dimensional structure diagrams generated with foRNA, (ii) compressed timeline representations illustrating structural transitions, and (iii) contact-frequency arc plots that reveal interaction persistence. These analyses include canonical Watson–Crick base pairs (A–U, G–C) and wobble base pairs (G–U). (i) ARNy Plotter generates dot-bracket notation from three-dimensional coordinates at each frame of the trajectory. This enables temporal tracking of base-pairing patterns and the identification of secondary-structure transitions across the ensemble. Dot–bracket notation provides a compact and interpretable representation of RNA secondary structure, facilitating both computational analysis and visualization. (ii) The generated dot-bracket strings are subsequently processed using foRNA (Force-directed RNA layout) [21] for interactive two-dimensional visualization. foRNA employs physics-based algorithms to produce aesthetically clear and structurally accurate secondary-structure diagrams that highlight base-pairing patterns and key structural motifs. (iii) Contact frequencies are computed for all nucleotide pairs across the entire trajectory, generating contact probability matrices that capture both persistent and transient interactions. Contact frequency data are visualized using arc plots, which depict interaction persistence across the trajectory and facilitate the identification of stable structural elements and dynamic interaction networks.

#### Interaction Networks

Understanding RNA tertiary structure requires detailed analysis of inter-nucleotide contacts beyond canonical Watson-Crick base pairs. ARNy Plotter implements comprehensive contact analysis algorithms that identify and quantify base pairing (canonical and non canonical) and stacking throughout the structural ensemble. For each structure, contacts are obtained from Barnaba annotations. Base pairing contact maps offer a detailed view of the RNA base-pairing network, classifying each contact according to the Leontis-Westhof scheme [22]. Stacking contact maps reveal nucleotide stacking interactions, distinguishing between parallel and antiparallel orientations. Both types of contact maps are dynamically updated as different structures are visualized in the 3D viewer. To help with compactness, the contacts represented in the map are between nucleotides.

#### Conformational landscape Analysis

Population histogram analysis provides crucial insights into the thermodynamic and kinetic properties of RNA conformational dynamics. ARNy Plotter generates projected landscapes by correlating structural order parameters such as RMSD or eRMSD with complementary metrics including native contact fraction (Q) and radius of gyration (R*_g_*). Given a reference structure, reference contact fraction Q is calculated as:

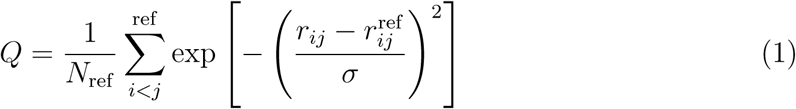

where *N_ref_*represents the number of contacts in the reference structure, *r_ij_* denotes the distance between residues *i* and *j*, and *σ* is a smoothing parameter.

Landscapes are presented as both 2D heatmaps using kernel density estimation and interactive 3D surfaces, enabling identification of metastable states, transition pathways, and conformational preferences. Clicking on a point on these maps will change the displayed frame to the closest one relative to the point selected, allowing to quickly discover what are the different states present in the ensemble.

#### Dimensional reduction analysis

Dimensionality reduction is crucial for interpreting high-dimensional structural ensembles from very large conformational ensembles. By projecting thousands of conformations into a low-dimensional space, these methods reveal dominant motions, highlight conformational clusters, and make complex dynamical landscapes easier to visualize and analyze. We have included in ARNy Plotter three different methods of dimensionality reduction: PCA, t-SNE and UMAP. PCA (Principal Component Analysis) identifies the directions of largest variance in the trajectory, providing a linear and interpretable decomposition of global motions. It excels at capturing large-scale collective movements but may overlook nonlinear relationships and fine structural distinctions. t-SNE (t-Distributed Stochastic Neighbor Embedding) focuses on preserving local similarities, making it effective at revealing discrete conformational clusters and subtle state separations. However, it distorts global distances, meaning it is less suited for understanding long-range structural relationships or continuous pathways. UMAP (Uniform Manifold Approximation and Projection) preserves both local and broader manifold structure, often capturing continuous transitions between states while still resolving distinct clusters. Compared with PCA and t-SNE, it provides a balance of global topology and local detail, making it useful for mapping complex conformational landscapes.

### System Architecture and Design Principles

ARNy Plotter employs a modern, scalable client-server architecture specifically designed to handle the computational demands of large-scale trajectory analysis while maintaining responsive user interactions. The system architecture separates computationally intensive analytical tasks from user interface operations, ensuring optimal performance across diverse usage scenarios. The platform integrates a combination of modern web technologies and Python-based computational tools to provide an interactive, scalable environment for ensemble analysis (Figure 2).

**Figure 2:**
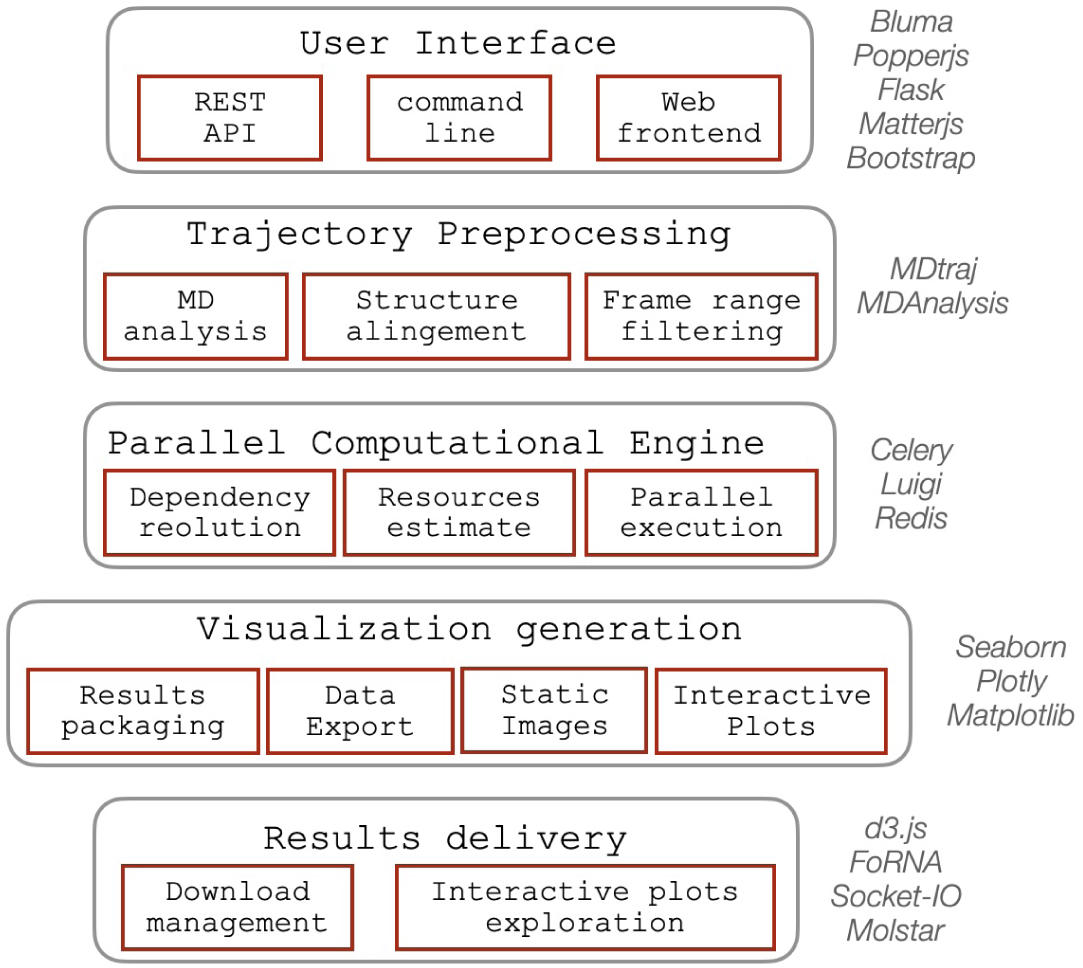
Schematic overview of the software’s backend architecture, illustrating the main functional components along with the external libraries and packages supporting each module.

Trajectory preprocessing relies on established libraries for molecular dynamics analysis, including MDAnalysis and MDTraj, which provide efficient routines for coordinate parsing, manipulation, operations and transcriptions to other formats. The computational backend is built around Flask, a lightweight Python web framework responsible for handling HTTP communication, managing user sessions, and rendering pages through Jinja2 templating. Analytical tasks are executed asynchronously through Celery, a distributed task-queue system that enables parallel processing of long-running computations. Celery provides fine-grained control over task execution, including error reporting, restart capabilities for failed jobs, and dynamic scaling with available computational resources. Communication between Celery workers and the web application is mediated by Redis, which handles message passing and stores intermediate results. Task dependencies are managed using Luigi, which constructs directed acyclic graphs of analytical steps. This ensures that shared intermediate computations, such as RMSD trajectories required by both conformational landscape analyses and RMSD-based visualizations, are performed once and propagated to downstream tasks, improving efficiency and speed. Real-time updates, including responsive recomputation of contact maps when users adjust trajectory frames or slider controls, are enabled through Socket.IO, which provides bidirectional communication between the client and server. The server layer is deployed using Gunicorn, which manages incoming user requests and distributes them across worker processes, ensuring high availability and scalability under concurrent usage.

The user interface combines Bootstrap and Bulma to deliver a modern, responsive layout compatible with a wide range of devices and screen resolutions. The command-line interface remains intentionally minimal, relying solely on plain Python to maximize accessibility for scripting and workflow integration. Interactive visualization is driven primarily by Plotly, which provides high-quality, browser-rendered graphics with native support for zooming, selection, and export. Additional specialized visualization libraries, including D3.js, FoRNA, and Matplotlib/Seaborn, are used for static graphics, RNA-specific structural depictions, and data summaries. For large trajectories containing thousands of points, the system leverages WebGL acceleration to offload rendering to the user’s graphics hardware when available, enabling fluid interactivity even for dense datasets. A custom results-delivery module integrates these visualization components into the front-end, using Plotly to maintain interactivity while ensuring consistent presentation and retrieval of computed results.

## 3 Results

We are now going to illustrate a few examples of applications of ARNy Plotter to structural ensembles of diverse origins.

The first ensemble consists of solution NMR structures of the P1–P2 frameshifting pseudoknot from the sugarcane yellow leaf virus (PDB ID: 2AP0) [23]. This ensemble contains 20 conformers generated by simulated annealing from random starting coordinates, followed by refinement using residual dipolar couplings. ARNy Plotter enables interactive visualization of the individual structures and facilitates comparison of their interaction networks through base-pair contact maps (Figure 3). These maps clearly reveal the main structural differences between conformers in terms of base pairing. All maps display a pseudoknot pattern, while frame-to-frame variations highlight specific interactions that appear or disappear. This analysis helps rationalize which structural features are robust, conserved across all conformers, and which elements are less certain, varying among individual models in the NMR ensemble.

**Figure 3:**
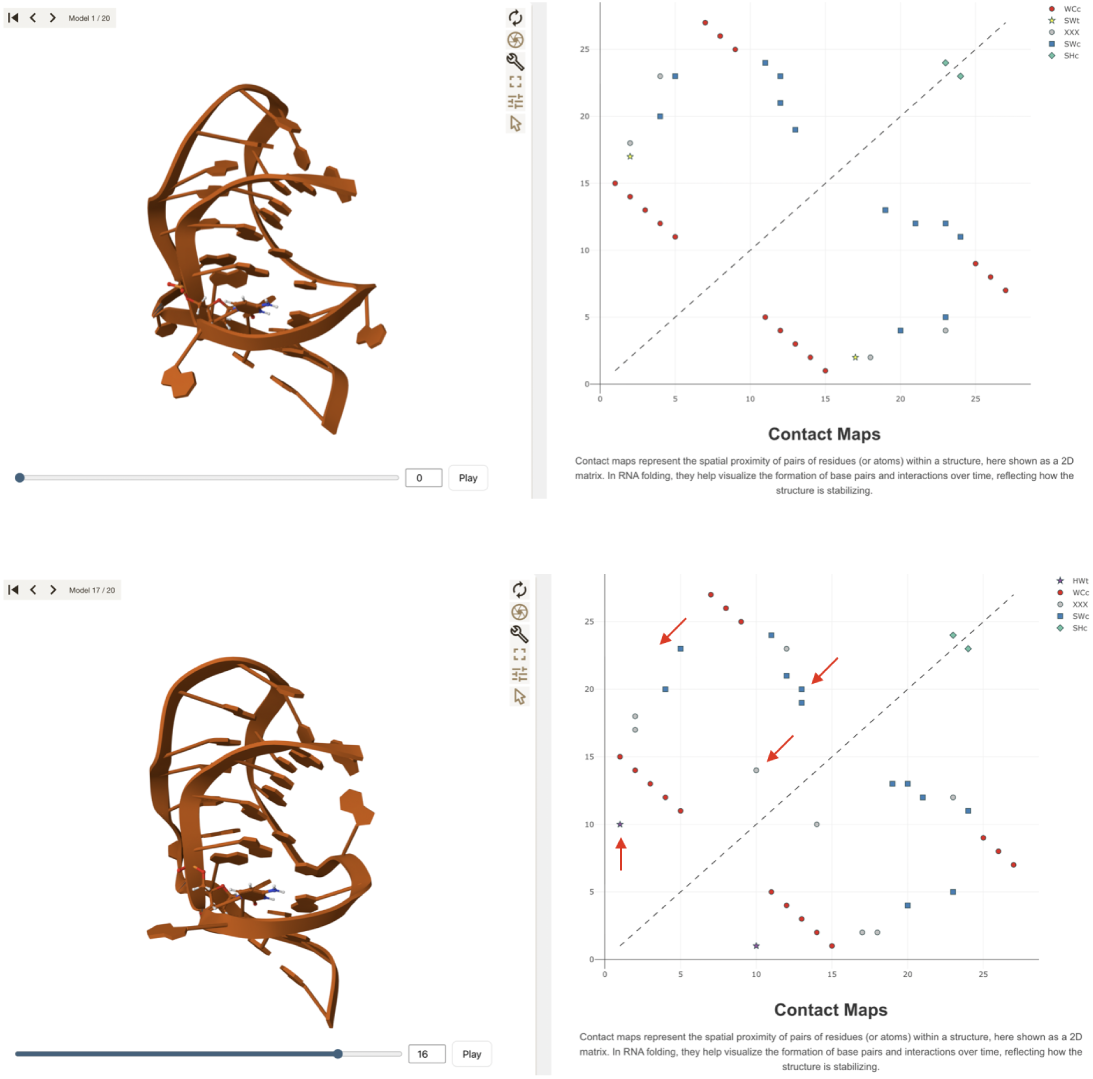
Visualization of two representative structures from the NMR ensemble of 2AP0. The left panels show the 3D conformations displayed in the interactive viewer, while the right panels present the corresponding base-pairing contact maps. Different symbols denote distinct interaction types according to the Leontis–Westhof classification. In the lower panels, red arrows have been added to highlight structural differences between the two ensemble members.

A second example is the analysis of 5300 structures of the Kaposi’s sarcoma-associated herpesvirus ORF50 transcript RNA (42 nucleotides), previously obtained through discrete path sampling simulations [24]. Previous analyses of the molecule’s energy landscape revealed two major energetic basins separated by a high energy barrier, each containing smaller sub-basins that capture structural variability while preserving the characteristic features of the parent basin (i.e., the same secondary structure). Using ARNy Plotter, we examined all structures identified as local minima through discrete path sampling (Figure 4). The PCA plot clearly highlights the partitioning of the ensemble into two principal basins, each comprising distinct subpopulations. Through the interactive functionality of the Result panel, clicking on a point in the PCA plot displays the corresponding structure along with all its associated features. As illustrated in the figure, this makes it straight-forward to identify key differences in both the 3D conformation and 2D organization of the molecule.

**Figure 4:**
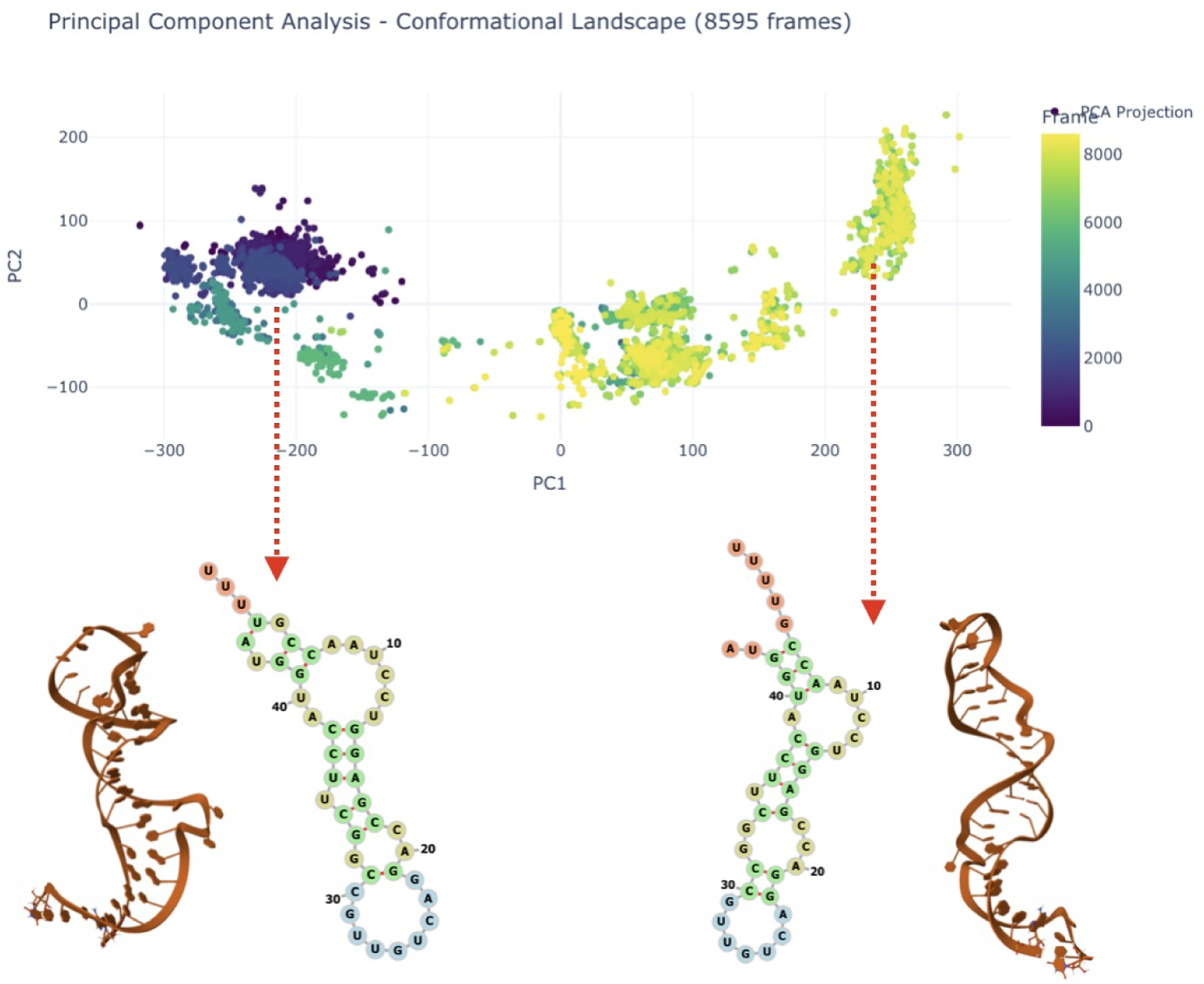
Analysis of the structural ensemble of ORF50 obtained from a discrete path sampling exploration of the molecule’s energy landscape. PCA reveals the partitioning of the ensemble in three main basins. Clicking on a point in the PCA plot allows to visualize the corresponding 3D structure and all its features (indicated here via the red arrow), including the 2D representation shown here.

The third example analyzes the ensemble of structures generated by a Hamiltonian Replica Exchange Molecular Dynamic simulation of the 5*^′^* hairpin of 7SK [25] (27 nucleotides), focusing specifically on those sampled under the unperturbed force field. Figure 5 illustrates the conformational landscape explored by the ensemble in which three distinct states are clearly visible when projecting over RMSD and fraction on native contacts, Q. By clicking on any point in the conformational landscape, ARNy Plotter automatically displays in the 3D viewer a representative structure corresponding to the selected (RMSD, Q) pair, and all structure-specific plots, such as contact maps, are updated accordingly. This interactive exploration makes it straightforward to retrieve structures directly from the landscape and to identify the key differences between conformational basins. In this specific example, the basin closest to the native structure (characterized by low RMSD and high Q values) preserves the triplet interaction observed in the crystal structure, formed by nucleotides 7, 20, and 24, which is known to be important for the interaction of 7SK with the protein HEXIM [26]. In contrast, the other two basins lose this feature and adopt conformations more similar to regular hairpins.

**Figure 5:**
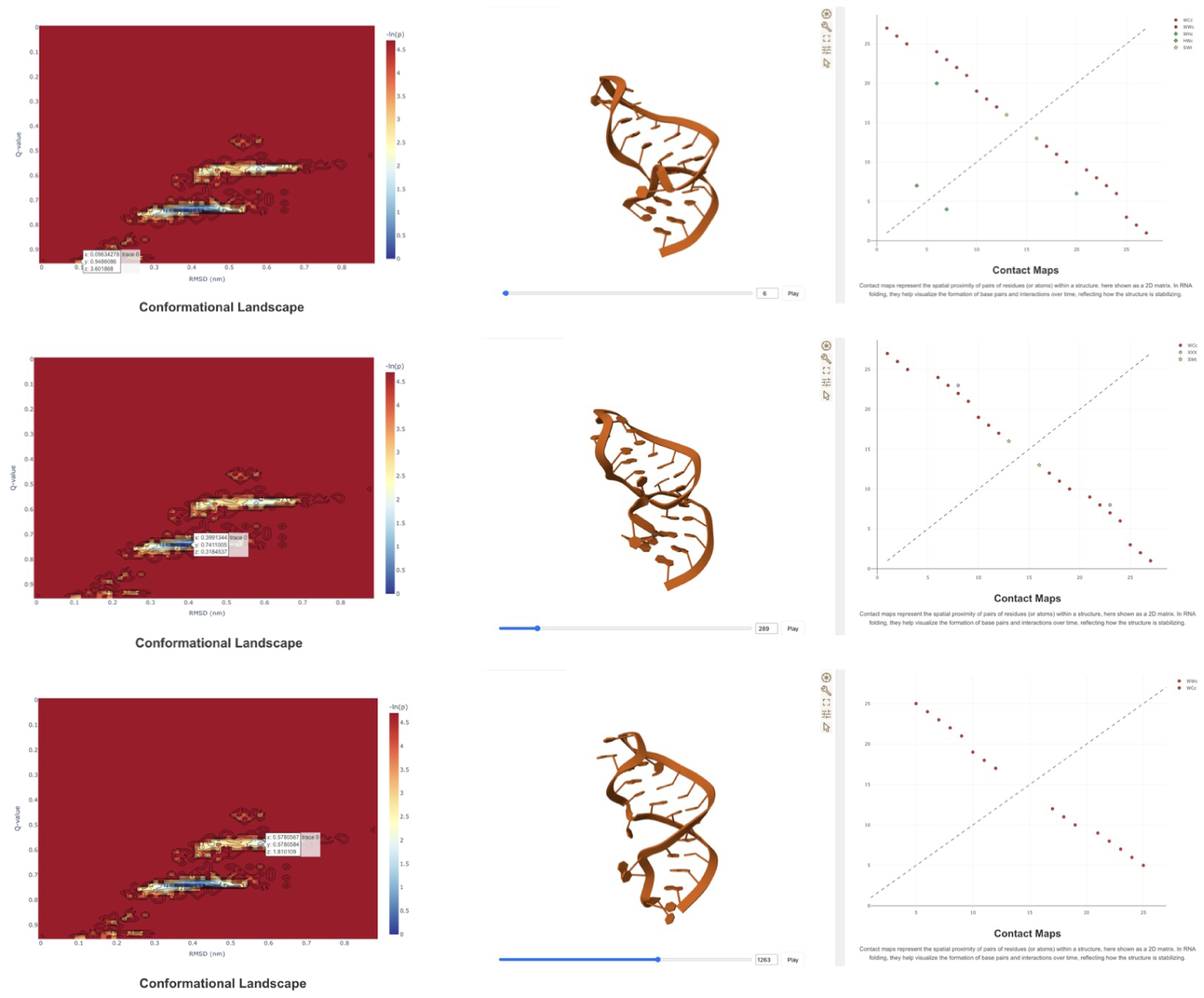
Conformational landscape analysis of the upper stem of 7SK highlighting 3 main structural basins. A representative of each basin can be selected by clicking on a point in the landscape (top, middle, bottom), allowing to show its 3D structure and detailed features, such as the contact map.

In the last example we use ARNy Plotter to analyze structures from a folding trajectory of the frameshifting pseudoknot of SARS-CoV-2 (76 nucleotides) starting from an unfolded structure, obtained through a ratchet MD simulation [27] biased toward the native pseudoknotted conformation (PDB id: 6XRZ [28]). ARNy Plotter analysis allows to clearly identify the folding pathway of the trajectory under several different metrics (Figure 6). The conformational landscape (6.C) reveals a narrow transition pathway in terms of RMSD and Q values. An intermediate state with a substantial population is observed at approximately 20 Å RMSD and a Q value of 0.5. Detailed inspection of this state indicates that it corresponds to the initial formation of the pseudoknot. The analysis of the successive formation of canonical base pairing (6.D) suggests that the pseudoknot forms early in the trajectory, and the arc-diagram, colored by contact frequency, highlights the persistence of one of the stems, which is almost always present (therefore forming early in the trajectory), while the other stem and the canonical pairs of the pseudoknot are less frequent, and stabilize toward the end of the trajectory (see (6.D)). Overall this analysis allows us to quickly understand the main features of the folding pathway for this specific trajectory.

**Figure 6:**
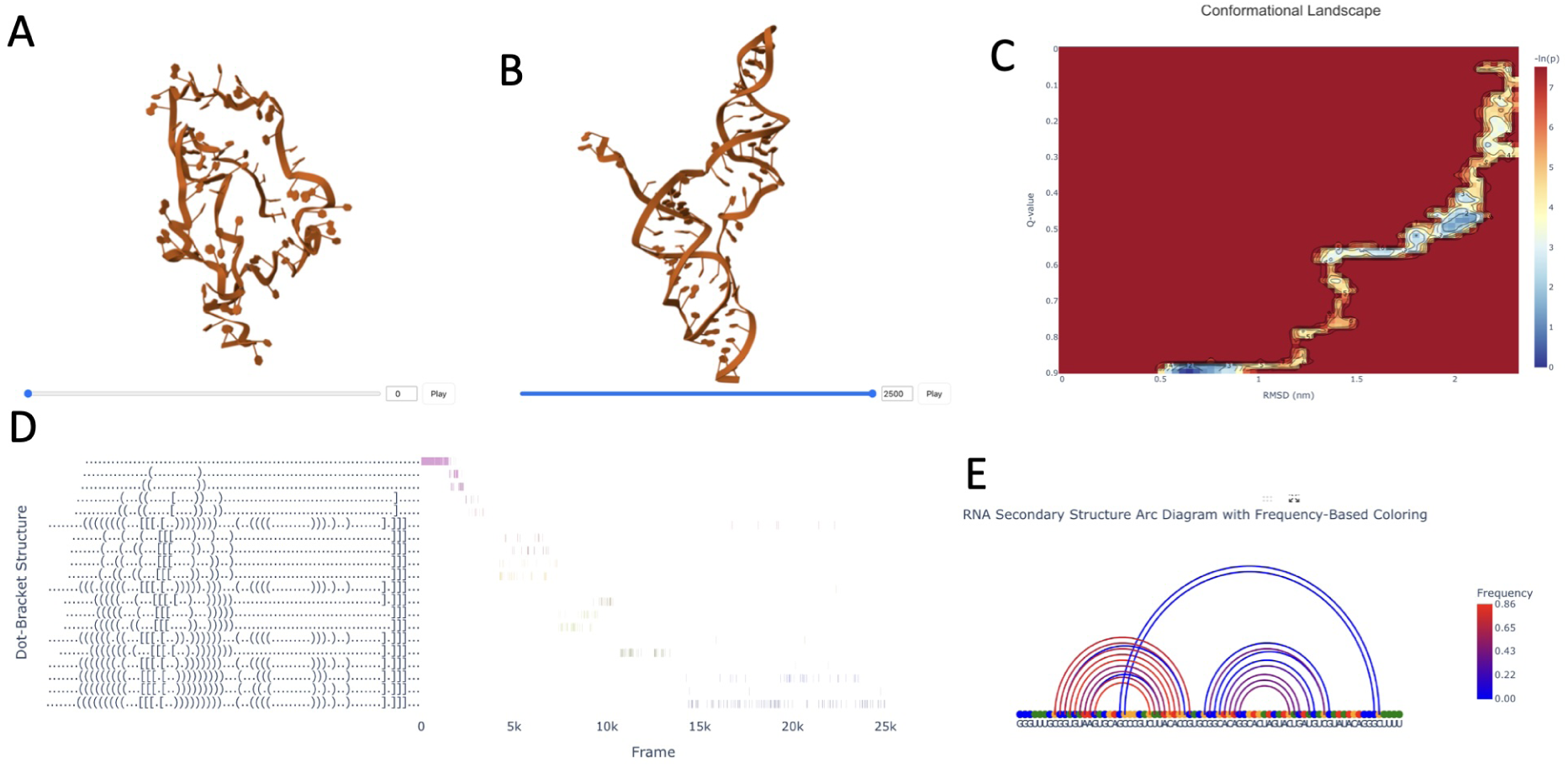
Results of the analysis of FSE rMD trajectory. A and B show the initial unfolded structure and the final target (native) structure respectively; C: conformational landscape of the folding process projected onto RMSD and Q-values with respect to target; D: time series of the formation of canonical base-pairs; E: arc-diagam the frequency of base pairs from which we can clearly observe the presence of the pseudoknot.

Having illustrated how ANRy Plotter captures key features of RNA structural ensembles through these examples, we now turn to a practical aspect that becomes critical when working with large data sets which is computing time. Efficient analysis is essential to keep such exploratory workflows usable, and ANRy Plotter addresses this challenge through optimized parallelization. In the reported examples, the maximum waiting time to obtain the complete analysis was approximately 2 minutes (Table 1), even for relatively large systems comprising thousands of structures. This limited processing time arises from the integrated design of the platform, in which all analyses are performed simultaneously. As a result, the trajectory is loaded only once, leading to a significant time savings. In addition, previously computed metrics (e.g., RMSD) are reused across multiple analyses and visualizations, further improving overall efficiency compared with running analyses individually.

**Table 1:** Performance benchmarks for ARNy Plotter ensemble analysis on representative RNA systems.

| System | Frames | Processing Time | Memory Usage |
| --- | --- | --- | --- |
| NMR structure 2AP0 | 19 | 600 ms | 2.6 MB |
| ORF50 path sampling | 5300 | 18 seconds | 184 MB |
| 7SK HREX | 40,000 | 121 seconds | 234 MB |
| FSE rMD | 10,000 | 120 seconds | 450 MB |

## 4 Availability and future directions

ARNy Plotter is freely accessible at https://arny-plotter.rpbs.univ-paris-diderot.fr/ and requires neither user registration nor software installation. The platform runs on all major operating systems using standard web browsers. Comprehensive documentation, step-by-step tutorials, and example datasets are provided to support new users. The complete source code is available at https://github.com/louismeuret/ARNy_Plotter under an open-source license, enabling community contributions as well as local installations for users with specific computational needs.

In future developments, we aim to further improve the computational efficiency of the platform. Although ARNy Plotter performs well for typical simulation sizes, extremely large trajectories or large systems can still require substantial processing time. However this limitation is less severe when using a local installation as jobs can run in the background, allowing users to still analyze at once very large trajectories and large systems.

As we continue developing tools for computing statistical interaction fields (SMIFS) around RNA molecules [29], aimed at detecting potential binding pockets and regions with preferred interaction propensities, we plan to integrate these analyses into future releases of ARNy Plotter. At present, such calculations can be performed on individual structures and short trajectories, but their computational and storage demands make them impractical for inclusion in the web server. While these features could eventually be supported in a local-installation version of the software, additional development work will be needed before full integration becomes feasible.

## 5 Conclusions

ARNy Plotter fills an important need in RNA structural biology by offering the first web-based tool designed specifically for analyzing and visualizing RNA structural ensembles. It brings together several useful analysis methods and modern visualization tools in one place, allowing researchers to explore complex RNA conformations without needing advanced computational skills. By unifying different types of RNA trajectory analysis, ARNy Plotter provides a clearer and more complete view of how RNA structures change and how these changes relate to function. This is especially important as RNA continues to gain attention for its many biological roles and its potential as a therapeutic target.

The platform offers several practical advantages. Its web interface removes installation and hardware barriers, making the tool easy for anyone to use. It also allows to easily share large structural ensembles and their analysis among collaborators, which is a key element to enhance the interplay between experimentalists and modelers. Interactive links between 3D structures and 2D plots help users connect visual and numerical information, and support for many file formats ensures compatibility with common simulation tools. Because of its ease of use and broad set of features, ARNy Plotter can support many kinds of RNA research, from basic studies of RNA folding to applications in drug design. By making advanced ensemble analysis accessible and straightforward, the platform can help more researchers work with RNA structure and speed up progress in the field.

## Acknowledgments

We thank the developers of MDAnalysis, Barnaba, foRNA, and MolStar for providing the foundational tools that enable ARNy Plotter functionality. We also acknowledge the beta users who provided valuable feedback during platform development.

We thank the RPBS platform for their support in the technical development and hosting of the web server.

